# Y chromosome and mitochondrial DNA haplogroups across behavioural traits in children from the general population

**DOI:** 10.1101/064444

**Authors:** Laurence J. Howe, A. Mesut Erzurumluoglu, George Davey Smith, Santiago Rodriguez, Evie Stergiakouli

## Abstract

**Objective:** To evaluate the association between Y chromosome and mitochondrial DNA haplogroups and a number of sexually-dimorphic behavioural and psychiatric traits.

**Methods:** The study sample included 4,211 males and 4,009 females with mitochondrial DNA haplogroups and 4,788 males with Y chromosome haplogroups who are part of the Avon Longitudinal Study of Parents and Children (ALSPAC). Different subsets of these populations were assessed using the Developmental and Well-being Assessment (DAWBA), Strengths and Difficulties Questionnaire (SDQ), SCDC (Social and Communication Disorder Checklist) and Psychotic Like Symptom Interview (PLIKSi) as measures of behavioural and psychiatric traits. Logistic regression was used to measure the association between haplogroups and the traits above.

**Results:** We found that the majority of behavioural traits in our cohort differed between males and females. However, Y chromosome and mitochondrial DNA major haplogroups were not associated with any of the variables. In addition, secondary analyses of Y chromosome and mitochondrial DNA subgroups also showed no association.

**Conclusion:** Y chromosome and mitochondrial DNA haplogroups are not associated with behavioural and psychiatric traits in a sample representative of the UK population.

## Introduction

Childhood psychiatric disorders are amongst the best examples of sexual dimorphism in disease. Males and females differ in terms of prevalence, age of onset, development and prognosis of psychiatric disorders.^1^ For example, the ratio of males to females in attention-deficit/hyperactivity disorder (ADHD) is between 2:1 and 8:1.^2^ However, research on possible factors contributing to sex differences has been limited.^3^ Many childhood psychiatric disorders, including ADHD and autism, can be conceptualized as extremes of continuous traits found in the general population with the same risk factors contributing to both clinical disease and subclinical traits. ^4-6^

One of the genetic factors that clearly differs between males and females is the sex chromosome complement. Y-linked genes are only expressed in males while some of the X-linked genes escape X chromosome inactivation and are expressed more highly in females. ^7^ In addition, some Y chromosome genes are expressed in the human brain ^7^ and the Y chromosome has been associated with increased aggression and impaired parental behaviour in animal models where the sex chromosome complement has been dissociated from the hormonal milieu.^8^ Patients with sex chromosome abnormalities have been found to be at increased risk of psychiatric disorders. There are reports of patients with Turner syndrome (XO) and Klinefelter syndrome (XXY) exhibiting higher rates of autism ^9^ while there is some evidence that individuals with 47, XYY syndrome have a higher risk of developing bipolar disorder and schizophrenia ^10^.

Despite its potential involvement in sexually dimorphic disorders and traits, the Y chromosome is largely excluded from genetic studies even of sexually dimorphic disease. The Y chromosome is mostly transmitted intact from father to son and is essential for sex differentiation through the SRY (Sex Determining Region of Y chromosome) ^11^. The male-specific region (MSY) of the chromosome does not recombine and is less than 1/6 of the size of the X chromosome retaining approximately only 17 genes ^11^. The lack of recombination provides the Y chromosome with the best haplotypic resolution in the human genome. However, the implication of lack of recombination is that the Y chromosome cannot be studied easily as part of a genome-wide association study (GWAS) pipeline. In contrast, it is analysed using haplogroups which are stable lineages of the Y chromosomes that share a common ancestor. The Y Chromosome Consortium (YCC) has produced a nomenclature system incorporating all verified Y chromosome SNPs and defining Y chromosome haplogroups ^12^. Studies have found evidence of associations between Y chromosome haplogroups and sexually dimorphic disease such as cardiovascular disease ^13^ while there are reports of Y chromosome micro deletions associated with infertility ^14^. However, Y chromosome haplogroups have not been found to be associated with autism ^15^.

Mitochondrial DNA is separate from nuclear DNA and is contained within mitochondria; intracellular organelles that provide energy to the cell. Its inheritance is analogous to the Y chromosome with DNA inherited unchanged, apart from mutations, from the mother and haplogroups used to study its association with disease ^16^. Studies have linked mitochondrial DNA with psychiatric disorders, predominantly schizophrenia ^17^ and bipolar disorder ^18^ with a review concluding that there is inconsistent evidence for the involvement of mitochondrial DNA in psychiatric disease ^19^.

The role of Y chromosome and mitochondrial DNA haplogroups in behavioural traits in children from the general population has not been investigated before. In this study we derived Y chromosome and mitochondrial DNA haplogroups in children from the Avon Longitudinal Study of Parents and Children (ALSPAC) and tested whether they are associated with parent-reported behavioural traits. We decided to focus on sexually dimorphic behavioural traits: Attention-Deficit/Hyperactivity Disorder (ADHD) traits, Oppositional Defiance Disorder (ODD) traits, Conduct Disorder (CD) traits, emotional traits, autistic traits, psychotic experiences and total behavioural traits.

## Methods

### Study Participants

The Avon Longitudinal Study of Parents and Children (ALSPAC) is a prospective birth cohort which recruited pregnant women with expected delivery dates between April 1991 and December 1992 from Bristol UK. 14,541 pregnant women were initially enrolled with 14,062 children born. Detailed information on health and development of children and their parents were collected from regular clinic visits and completion of questionnaires. A detailed description of the cohort has been published previously ^20 21^. The study website contains details of all the data that is available through a fully searchable data dictionary: http://www.bris.ac.uk/alspac/researchers/data-access/data-dictionary/. Ethical approval was obtained from the ALSPAC Law and Ethics Committee and the Local Ethics Committees.

## Genotyping of Y chromosome and mitochondrial DNA

A total of 9,912 participants were genotyped using the Illumina HumanHap550 quad genome-wide SNP genotyping platform by Sample Logistics and Genotyping Facilities at the Wellcome Trust Sanger Institute and LabCorp (Laboratory Corporation of America) using support from 23andMe. PLINK software (v1.07) was used to carry out quality control (QC) measures ^22^. Individuals were excluded from further analysis on the basis of having incorrect gender assignments, minimal or excessive heterozygosity (< 0.320 and > 0.345 for the Sanger data and < 0.310 and > 0.330 for the LabCorp data), disproportionate levels of individual missingness (> 3%), evidence of cryptic relatedness (> 10% IBD) and being of non-European ancestry (as detected by a multidimensional scaling analysis seeded with HapMap 2 individuals). EIGENSTRAT analysis revealed no additional obvious population stratification and genome-wide analyses with other phenotypes indicate a low lambda) ^23^. SNPs with a minor allele frequency of < 1% and call rate of < 95% were removed. After QC, 8,365 unrelated individuals were available for analysis.

For Y-DNA haplogroup determination, the Y-chromosomal SNPs of all 5,085 male participants in the dataset were used. The pseudo-autosomal SNPs (coded as chromosome 25) were removed from the analysis using the PLINK software package ^22^. The resulting Y chromosomal genotypes (816 SNPs) of each individual were then piped in to the Y-Fitter (v0.2) software (maps genotype data to the Y-DNA phylogenetic tree built by Karafet *et al* ^24^, available online at sourceforge.net/projects/yfitter) and their respective Y-DNA haplogroup was determined. After removal of individuals with ‘False’ haplogroup determinations (i.e. ones which did not have enough SNPs to reliably determine haplogroup), we were left with 5,080 individuals. Remaining individuals with a haplogroup result which began with the letter R (e.g. R1b1) were clustered in to a single group named ‘Clade R’ and likewise the same was done with the haplogroups beginning with the other letters. Major haplogroups were defined as containing multiple haplogroups that are closely related to utilize information on less common haplogroups. After QC and removing individuals with withdrawn consent; our data-set contained 4,788 males with derived Y chromosome haplogroups.

7,554 custom mitochondrial probes, targeting 2,824 unique mtDNA positions, were included on the Illumina HumanHap550 quad genome-wide SNP genotyping platform. All heterozygous genotype calls (i.e. heteroplasmy) were set to missing prior to quality control using PLINK (Purcell, Neale *et al*. 2007). Genotype calls obtained from each probe were compared to the human mitochondrial database of non-pathological mitochondrial sequence variants (www.hmtdb.uniba.it:8080/hmdb/) to ensure that known allelic variants were being called. Probes were excluded in cases where genotype calls were not represented in the Cambridge Reference Sequence reference (rCRS) or one of the known allelic variants. Probes with an overall call rate of < 95% were excluded prior to analysis. The genotyping concordance of the remaining probes was investigated by comparing the genotype calls in 445 replicate samples. With the exception of probe failure (i.e. missing data), a 100% genotyping concordance rate was obtained for each probe. All probes with a failure rate of > 5% in the replicate sample were further excluded. In cases where multiple probes passed the above mentioned QC criteria, the probe with the highest calling rate was used for analysis. A total of 1062 probes passed QC for the batch that was genotyped by Laboratory Corporation of America (n=7590), whilst 629 probes passed QC for the batch genotyped by the Sanger Institute (n=775). Haplogroup assignment was performed as described by Kloss-Brandstatter et al ^25^, and samples with a quality score of more than 90% were used for our analysis. Major haplogroups were defined as containing multiple haplogroups that are closely related to utilize information on less common haplogroups. After QC and removing individuals with withdrawn consent; our data-set contained 4,211 males and 4,009 females with derived mitochondrial DNA haplogroups.

## Behavioural Trait Variables

The Strengths and Difficulties Questionnaire (SDQ) is a behavioural screening questionnaire with high reliability and validity which includes questions on five domains: emotional symptoms, conduct problems, hyperactivity symptoms, peer relationship problems and prosocial behaviours ranging from 0 to 10 each ^26^. A total difficulties score can be obtained by summing the following subscales: emotional symptoms, conduct problems, hyperactivity score and peer relationship problems (not including prosocial behaviour for which a higher score indicates less behavioural problems). SDQ total difficulties score ranges from 0 to 40 (www.sdqinfo.com). Mother’s reports on their children’s behavioural problems were obtained at 7 years of age. Cut-offs were used to dichotomise each of the five SDQ subscales as well as total difficulties score according to previous studies.

The Development and Well-Being Assessment (DAWBA) is a semi-structured interview used to diagnose psychiatric disorders in children ^27^ and calculate the number of psychiatric disorder traits. Attention-Deficit/Hyperactivity Disorder (ADHD), Oppositional/Defiant Disorder (ODD) and Conduct Disorder (CD) traits were assessed in ALSPAC when the participants were aged 7 years and 7 months old using the parent-completed DAWBA attention/activity symptoms score, awkward behaviours score and troublesome behaviours score respectively. For each item, parents marked boxes to say whether their child showed the behaviour; these were coded 0 for “no,” 1 for “a little more than others”, and 2 for “a lot more than others.” A total trait score was calculated by summing these responses. Scores on measures with <30% missing items were mean imputed.

The Skuse social cognition scale (SCDC) ^28^ is designed to summarise the main features of the social cognition behaviour of individuals and was employed to assess autistic traits in children from ALSPAC. This was in the form of a parent-completed screening questionnaire on the child at 7 years and 7 months.

The semi-structured Psychosis-Like Symptom Interview (PLIKSi) ^29,30^, which draws on principles of standardized clinical examination developed for the Schedule for Clinical Assessment in Neuropsychiatry (SCAN), was used to assess psychotic experiences at ages 12 and 18 years. The PLIKSi allows rating of 12 psychotic experiences including hallucinations (visual and auditory), delusions (spied on, persecution, thoughts read, reference, control, grandiosity, other) and experiences of thought interference (broadcasting, insertion and withdrawal). The interviewers were psychology graduates trained in assessment using the SCAN psychosis section and using the PLIKSi. Psychotic experiences were rated as not present, suspected or definitely psychotic. Unclear responses were always ‘rated down’ and symptoms were rated as definite only when a clear example was provided. Individuals were deemed as having a psychotic experience if rated as having 1 or more definite or suspected psychotic experience compared to no psychotic experiences.

### Statistical Analysis

Most of the continuous variables were heavily zero-skewed; the majority of participants scored zero. We used categorical variables as appropriate normalisation was not possible in the majority of cases. Cut-offs were used to dichotomise each of the variables according to previous studies and the distribution of the variables ^26^. The attention/activity symptoms score and SCDC were stratified into (≤5 & >5), the awkward behaviours score, troublesome behaviours score and two PLIKS measures were stratified into (≤0 & >0), the hyperactivity traits score and emotional symptoms score were stratified into (≤2 & >2), the conduct traits score was stratified into (≤1 & >1) and the total behavioural score was stratified into (≤10 & >10).

Chi squared tests were used to test for sex differences in categorical variables with linear regression used for continuous variables. Multiple regression was used to test for association between haplogroups and continuous variables whilst adjusting for confounders. Chi squared tests were used to test for associations between haplogroups and categorical variables with logistic regression employed when confounders were included. The correlation between the continuous variables was computed to determine if the variables were independent with a 0.8 cut-off. The Bonferroni-Holm method was used to adjust the number of independent variables for multiple testing ^31^.

The pattern of inheritance means that individuals with different haplogroups may differ phenotypically, specifically in terms of social class and outcomes associated with it, such as education. The most appropriate variables to gauge social class of a child would be the social class and the educational level of the parents. General certificate of secondary education (GCSE) results were used as a measure of a child’s cognitive abilities. The Y chromosome is inherited from the father; therefore paternal social class was included in the Y chromosome analysis. Mitochondrial DNA is inherited from the mother; thus maternal social class was included in the analysis along with paternal social class for mitochondrial DNA. Y chromosome analysis was only performed on male participants while male and female participants were included in mitochondrial DNA analysis.

Initially just the major haplogroups e.g. “R” and “I” for the Y chromosome and “H” and “U” for the mitochondrial DNA were analysed but subsequent analysis investigated subgroups of these major haplogroups. Subgroups were investigated if their frequency was >2%. Y chromosome and mitochondrial DNA haplogroups were grouped together using online phylogenetic trees ^32^ ^33^.

Pie-charts were created using ALSPAC haplogroup data as well as haplogroup data for England from Eupedia (www.eupedia.com) and STATA 13 was used for all data analysis.

## Results

Y chromosome and mitochondrial DNA haplogroups were derived from children in the general population. The most common major Y chromosome haplogroups in ALSPAC boys were R (72%) and I (19%) Y-chromosome haplogroups. The distribution of major Y chromosome haplogroups in ALSPAC was consistent with available data from England (**Figure 1**). The most common major mitochondrial DNA haplogroups in ALSPAC were H (49%), U (13%), J (11%) and T (10%). The distribution of the major mitochondrial-DNA haplogroups in ALSPAC was consistent with available data from England (**Figure 1**).

**Figure 1.**
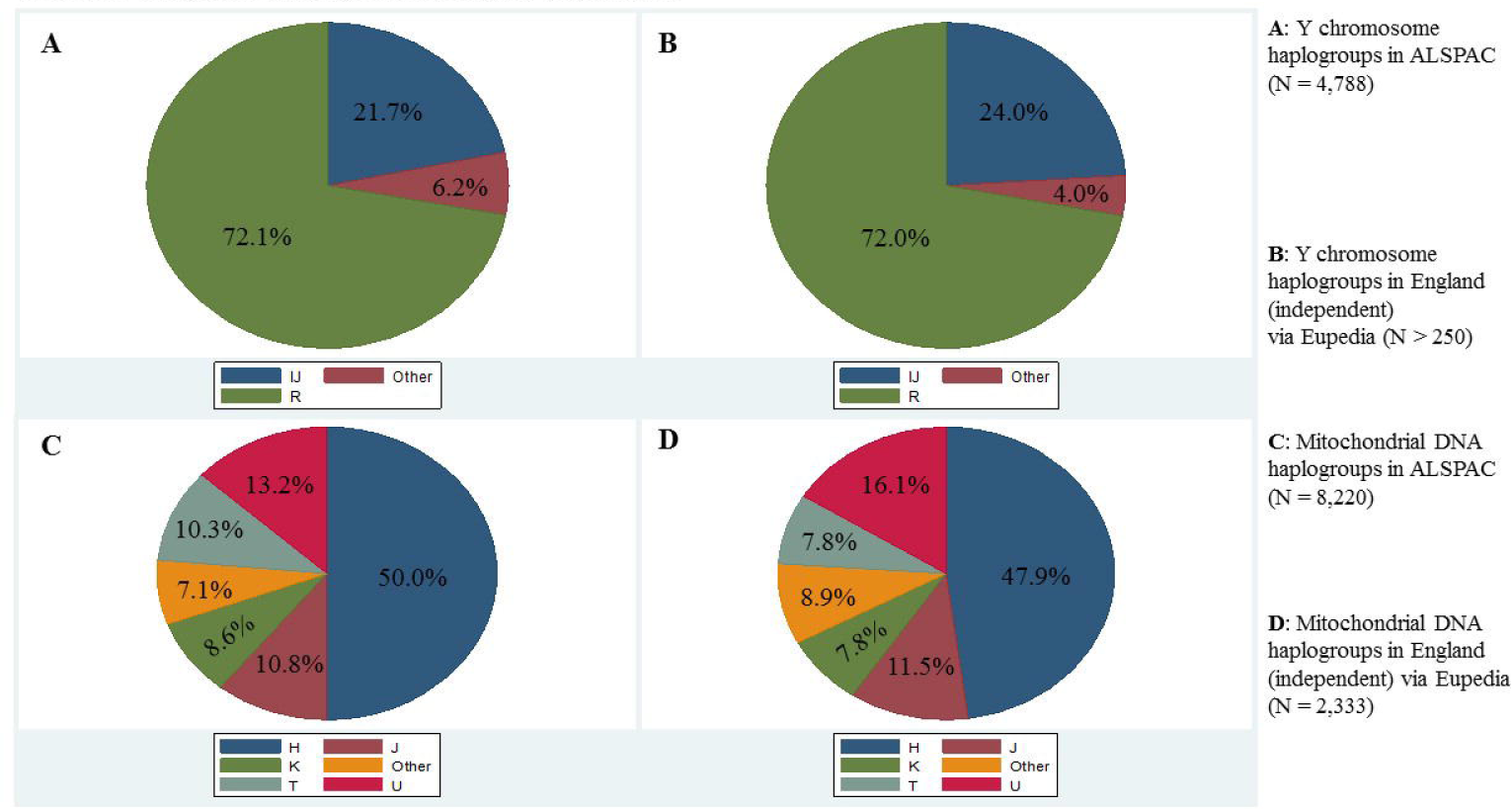
Y chromosome and Mitochondrial DNA haplogroups in ALSPAC compared to independent English populations

We investigated whether behavioural and psychiatric traits differed between males and females. The majority of traits showed sex differences with the exception of psychosis measured at 14 (**Figure 2**); there was a higher proportion of males than of females with high number of behavioural and psychiatric traits across all categories. There was one exception, psychotic experiences, which were more commonly reported in females. The largest difference was for attention/activity trait, with 38.1% of males scoring high compared to 24.5% of females (P < 0.001).

**Figure 2.**
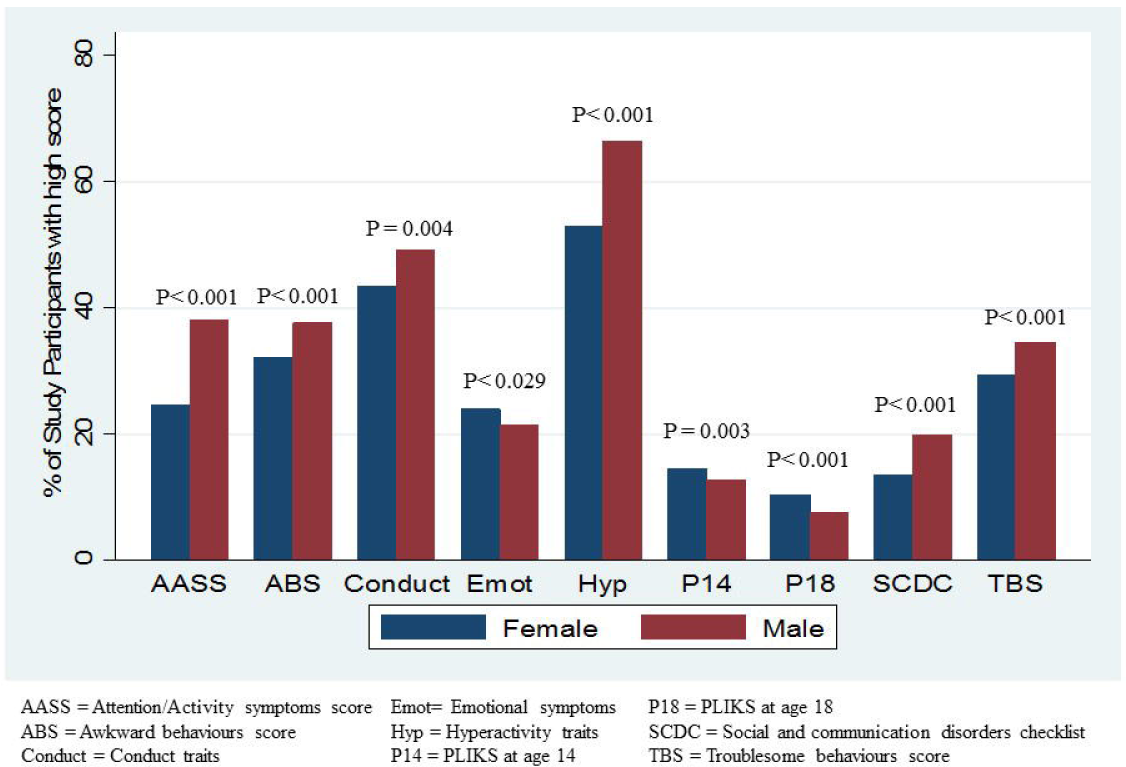
Sex differences in psychiatric trait scores (N > 4000)

The primary analysis involved Y chromosome and mitochondrial DNA major haplogroups. Tetrachoric correlation was computed between the 10 psychiatric trait binary variables, no pairwise correlation exceeded the threshold of 0.8 (**Table S1**). Therefore multiple testing correlation was performed assuming 10 independent variables. Major Y chromosome haplogroups were not associated with a high or low classification of number of behavioural and psychiatric traits (**Table 1**) or the raw number of traits (**Table S2**).

**Table 1.** Odds ratios of major Y chromosome haplogroups on binary behavioural trait measures from logistic regression adjusted for paternal social class and GCSE results.

| Behavioural/ Psychiatric trait score | Odds ratios of major Y chr haplogroup on high psychiatric trait category: OR (95% C.I.)<br>R haplogroup is the reference |  | Adjusted P values |
| --- | --- | --- | --- |
|  | I | Other |  |
| Attention/ Activity symptoms score (DAWBA)<br>N=2400 | 1.08 (0.88, 1.33) | 1.46 (1.02, 2.09) | 0.10 |
| Awkward behaviours score (DAWBA)<br>N=2390 | 1.06 (0.86, 1.30) | 1.31 (0.92, 1.86) | 0.31 |
| Troublesome behaviours score (DAWBA)<br>N=2396 | 0.98 (0.79, 1.20) | 1.03 (0.72, 1.49) | 0.95 |
| SCDC<br>N=2397 | 0.98 (0.76, 1.27) | 1.28 (0.84, 1.96) | 0.50 |
| **Hyperactivity traits (SDQ)**<br>**N=2461** | 1.06 (0.86, 1.31) | 1.12 (0.78, 1.62) | 0.74 |
| **Conduct traits (SDQ)**<br>**N=2462** | 1.34 (1.10, 1.63) | 1.11 (0.78, 1.56) | 0.014 > 0.006<sup>1</sup> |
| **Emotional symptoms (SDQ)**<br>**N=2462** | 1.05 (0.83, 1.33) | 1.20 (0.81, 1.79) | 0.64 |
| **Total behavioural traits (SDQ)**<br>**N=2459** | 1.01 (0.80, 1.27) | 1.30 (0.89, 1.89) | 0.41 |
| **PLIKSi age 14**<br>**N=2062** | 0.89 (0.64, 1.22) | 0.94 (0.53, 1.65) | 0.76 |
| **PLIKSi age 18**<br>**N=1270** | 0.99 (0.59, 1.67) | 1.20 (0.50, 2.90) | 0.92 |

95% CI: 95% Confidence Interval; DAWBA: Development and Well-Being Assessment; SCDC: Social and communication disorders checklist; SDQ: Strengths and Difficulties Questionnaire; PLIKSi: Psychosis-Like Symptom Interview
1 Smallest adjusted p value compared to multiple testing threshold

Weak evidence was found for association between the major mitochondrial-DNA haplogroups and a high or low classification of number of behavioural and psychiatric traits (**Table 2**) or the raw number of traits (**Table S3**).

**Table 2.** Odds ratios of major Mitochondrial DNA haplogroups on binary behavioural trait measures from logistic regression adjusted for maternal and paternal social classes and GCSE results.

| Odds ratios of major mitochondrial DNA haplogroup on high psychiatric trait category: OR (95% C.I.)<br>HV haplogroup is the reference |  |  |  |  |  | Adjusted P values |
| --- | --- | --- | --- | --- | --- | --- |
| Behavioural/<br>Psychiatric trait score | J | K | TR | U | Other |  |
| **Attention/ Activity symptoms score (DAWBA)**<br>**N=3857** | 1.02<br>(0.80, 1.29) | 0.84<br>(0.64, 1.10) | 1.04<br>(0.82, 1.31) | 1.00<br>(0.80, 1.25) | 0.86<br>(0.65, 1.15) | 0.72 |
| **Awkward behaviours score (DAWBA)**<br>**N=3841** | 0.93<br>(0.74, 1.17) | 0.92<br>(0.72, 1.18) | 0.90<br>(0.72, 1.13) | 0.97<br>(0.78, 1.20) | 0.84<br>(0.64, 1.10) | 0.79 |
| **Troublesome behaviours score (DAWBA)**<br>**N=3843** | 0.88<br>(0.69, 1.11) | 0.99<br>(0.76, 1.28) | 0.99<br>(0.79, 1.25) | 1.03<br>(0.83, 1.27) | 1.15<br>(0.88, 1.51) | 0.74 |
| **SCDC**<br>**N=3850** | 0.86<br>(0.63, 1.16) | 0.84<br>(0.60, 1.18) | 0.86<br>(0.64, 1.16) | 0.96<br>(0.73, 1.26) | 0.76<br>(0.52, 1.09) | 0.59 |
| **Hyperactivity traits (SDQ)** | 0.89<br>(0.71, 1.11) | 1.02<br>(0.80, 1.30) | 1.08<br>(0.87, 1.35) | 0.93<br>(0.75, 1.14) | 1.07<br>(0.82, 1.38) | 0.73 |

| **N=3926** |  |  |  |  |  |  |
| --- | --- | --- | --- | --- | --- | --- |
| **Conduct traits (SDQ)**<br>**N=3925** | 1.21<br>(0.98, 1.50) | 0.89<br>(0.70, 1.12) | 1.02<br>(0.82, 1.26) | 0.93<br>(0.76, 1.13) | 0.92<br>(0.71, 1.18) | 0.30 |
| **Emotional symptoms (SDQ)**<br>**N=3927** | 1.10<br>(0.87, 1.38) | 0.84<br>(0.64, 1.11) | 0.92<br>(0.72, 1.17) | 0.95<br>(0.76, 1.19) | 0.79<br>(0.59, 1.06) | 0.52 |
| **Total behavioural traits (SDQ)**<br>**N=3922** | 1.26<br>(0.98, 1.63) | 1.11<br>(0.83, 1.47) | 1.19<br>(0.92, 1.54) | 1.11<br>(0.87, 1.41) | 0.77<br>(0.55, 1.08) | 0.14 |
| **PLIKSi age 14**<br>**N=3426** | 1.09<br>(0.78, 1.51) | 0.88<br>(0.61, 1.27) | 1.14<br>(0.84, 1.56) | 0.70<br>(0.50, 0.98) | 0.80<br>(0.53, 1.22) | 0.19 |
| **PLIKSi age 18**<br>**N=2369** | 0.95<br>(0.57, 1.59) | 1.07<br>(0.64, 1.78) | 1.65<br>(1.09, 2.50) | 1.15<br>(0.75, 1.79) | 0.47<br>(0.21, 1.03) | 0.05 > 0.006<sup>1</sup> |

95% CI: 95% Confidence Interval; DAWBA: Development and Well-Being Assessment; SCDC: Social and communication disorders checklist; SDQ: Strengths and Difficulties Questionnaire; PLIKSi: Psychosis-Like Symptom Interview <sup>1</sup> Smallest adjusted p value compared to multiple testing threshold

To further investigate for potential associations, we considered subgroups of the major Y chromosome haplogroups R and I (**Table 3)** in the secondary analysis. Y chromosome haplogroups and subgroups were not associated with categorical (**Table S4**) or continuous variables (**Table S5**) of behavioural and psychiatric traits.

We considered subgroups of the larger mitochondrial-DNA haplogroups H, J, K, T and U (**Table 3**) in the secondary analysis. Weak evidence was found of association between these haplogroups and subgroups and categorical (**Table S6**) or continuous variables (**Table S7)** of behavioural and psychiatric scores.

**Table 3.** Major haplogroups and subgroups.

| **Y chromosome** |  | **Mitochondrial DNA** |  |
| --- | --- | --- | --- |
| Haplogroup | Count (%) | Haplogroup | Count (%) |
| **R** | **3452 (72.1)** | **H** | **4109 (49.9)** |
| R1a1 | 210 (4.4) | H1 | 1592 (19.4) |
| R1b1 | 328 (6.9) | H2 | 826 (10.0) |
| R1b1b2 | 1528 (31.9) | H3 | 387 (4.7) |
| R1b1b2g | 1061 (22.2) | H4 | 168 (2.0) |
| R1b1b2h | 296 (6.2) | H5 | 252 (3.1) |
| Q and other R | 29 (0.6) | H6 | 235 (2.9) |
| **IJ** | **1039 (21.7)** | Other HV | 649 (7.8) |
| I1 | 460 (9.6) | **J** | **885 (10.8)** |
| I2 | 150 (3.1) | J1 | 717 (8.7) |
| I2b | 228 (4.8) | J2 | 168 (2.1) |
| Other I | 71 (1.5) | **K** | **706 (8.6)** |
| J | 130 (2.7) | K1 | 569 (6.9) |
| **Other** | **297 (6.2)** | Other K | 137 (1.7) |
| E | 143 (3.0) | **T** | **845 (10.3)** |
| CFGH | 110 (2.3) | T2 | 668 (8.1) |
| LTNO | 44 (0.9) | Other TR | 177 (2.2) |
|  |  | **U** | **1084 (13.2)** |
|  |  | U4 | 165 (2.0) |
|  |  | U5 | 722 (8.8) |
|  |  | Other U | 197 (2.4) |
|  |  | **Other** | **591 (7.2)** |
|  |  | M (M, C, D, L) | 67 (0.8) |
|  |  | N (N, A, I, W, X) | 524 (6.4) |
| Total: | **4788** | **Total:** | **8220** |

## Discussion

Research on possible factors contributing to sex differences in behavioural and psychiatric traits has been limited ^3^ and this is the first investigation of the association of Y chromosome and/or mitochondrial DNA haplogroups with behavioural and psychiatric traits in children from the general population. We report here that there were more boys than girls with higher scores on behavioural and psychiatric trait scales. However, psychotic experiences were more common in girls than boys at two different time points. The haplogroup structure of Y chromosome and mitochondrial DNA in ALSPAC is consistent with a population of English origin. Y chromosome or mitochondrial DNA haplogroups were not associated with behavioural and psychiatric traits in ALSPAC.

There is convincing evidence that the Y chromosome is implicated in regulating differences in behaviour between the sexes in animal models. However, the biological mechanisms behind this are poorly understood. The difficulty in including the Y chromosome in GWAS is not contributing to improving understanding of the role of Y chromosome. The most convincing report of an association of the Y chromosome with a sexually dimorphic phenotype has been found in relation to coronary heart disease. In a cohort of British men haplogroup I was associated with a 50% increase in risk of cardiovascular disease with differential expression of macrophages between haplogroup I and other haplogroups ^13^.

Behavioural problems and psychiatric disorders in children and adults are highly sexually dimorphic. However, there are few reports of Y chromosome haplogroup investigations in psychiatric phenotypes. A previous study ^15^ found weak evidence of an association between Y chromosome haplogroups and autism and our findings are consistent with this. In a study of Y chromosome haplogroups and aggression in a cohort of men from Pakistan, there were no differences in mean scores of aggression across the five different haplogroups captured in the study ^34^. Mitochondrial DNA has been investigated in relation to psychiatric disorders, although conclusions have been limited with small sample sizes and inconsistent reporting ^19^.

We investigated Y chromosome and mitochondrial DNA haplogroups in children from a sample representative of the general population. The main strength of our study was the use of a large population of children from a homogeneous population. The ALSPAC study participants come from a small geographic region in the South West of England. Only participants of European origin were included in the study as part of the quality control protocol of the GWAS. All of our analyses were performed within the same population and did not require a control group, which could have resulted in issues with population stratification due to imperfect matching of cases and controls. This can be an important issue in Y chromosome analysis because even populations from the same country can have different Y chromosome haplogroup structure. We were also able to access data on different behavioural and psychiatric trait scales, such as the SDQ and DAWBA, which have been used extensively in population cohorts for research purposes.

One of the limitations of our study was the small number of individuals in some of the haplogroups. However, this is expected due to the rarity of certain Y chromosome haplogroups in European populations. Another challenge in the study was presented by the distribution of behavioural and psychiatric traits in the general population. A large number of individuals did not have any symptoms. Therefore, the distribution of the traits was exceedingly left skewed and transforming to something approaching normality was not possible. To overcome this, we dichotomized the variables and also performed sensitivity analyses with raw continuous scores. Also, we only investigated common variants and not copy-number variation or rare variants.

Although we do not report an association between the Y chromosome haplogroups and mitochondrial DNA haplogroups and risk of behavioural and psychiatric problems in the general population, we consider that our study has important implications; it highlights that if there are common variants on the Y chromosome and mitochondrial DNA associated with behavioural problems, they are of small effect size and would require large collaborative efforts to be identified. Given that the DNA markers on the Y chromosome and mitochondrial DNA are available on most GWAS chips but have not been largely utilised, such a collaborative effort would involve extracting relevant data from multiple studies rather than new genotyping efforts and could be an important step in understanding non-autosomal variation. Collaboration would be even more important for the Y chromosome because only half of any genotyped sample is usually male and can be used for analysis.

## Acknowledgements

This work was supported by the Medical Research Council and the University of Bristol [MRC Integrative Epidemiology Unit (IEU) MC_UU_12013/1-9].

The authors are extremely grateful to all the families who took part in the ALSPAC study, the midwives for their help in recruiting them, and the whole ALSPAC team, which includes interviewers, computer and laboratory technicians, clerical workers, research scientists, volunteers, managers, receptionists, and nurses. ALSPAC GWAS data was generated by Sample Logistics and Genotyping Facilities at the Wellcome Trust Sanger Institute and LabCorp (Laboratory Corporation of America) supported by 23andMe. We thank Drs. David Evans, Beate St Pourcain, Susan Ring, John Kemp and George McMahon for their contribution to QC of ALSPAC GWAS data. We especially thank Dr. John Kemp for his contribution to the QC of Y chr genetic data.

