## Supplementary Materials for "Y chromosome and mitochondrial DNA haplogroups across behavioural traits in children from the general population"

**Supplementary Material**

**Table S1. Pairwise tetrachoric correlations between binary behavioural trait variables**

**Table S2. Association of major Y chromosome haplogroups in ALSPAC with the number of** **behavioural traits adjusted for paternal social class and GCSE results**

**Table S3: Association of major mitochondrial DNA haplogroups in ALSPAC with the number of behavioural traits adjusted for maternal and paternal social class and GCSE results**

**Table S4. Odds ratios of Y chromosome haplogroups subgroups on binary psychiatric trait measures from logistic regression adjusted for paternal social class and GCSE results**

**Table S5. Association of Y chromosome haplogroup subgroups in ALSPAC with the number of behavioural traits adjusted for paternal social class and GCSE results**

**Table S6. Odds ratios of mitochondrial DNA chromosome haplogroup subgroups on binary behavioural trait measures from logistic regression adjusted for paternal social class and GCSE results**

**Table S7. Association of mitochondrial DNA haplogroup subgroups in ALSPAC with the number of behavioural traits adjusted for maternal and paternal social classes and GCSE results**

**Table S1. Pairwise tetrachoric correlations between binary behavioural trait variables**

|  | AASS | ABS | TBS | SCDC | Hyper | Conduct | Emotional | Total | PLIKS 14 | PLIKS 18 |
| --- | --- | --- | --- | --- | --- | --- | --- | --- | --- | --- |
| AASS | 1 | - | - | - | - | - | - | - | - | - |
| ABS | 0.57 | 1 | - | - | - | - | - | - | - | - |
| TBS | 0.39 | 0.33 | 1 | - | - | - | - | - | - | - |
| SCDC | 0.67 | 0.67 | 0.48 | 1 | - | - | - | - | - | - |
| Hyper | 0.72 | 0.37 | 0.33 | 0.48 | 1 | - | - | - | - | - |
| Conduct | 0.48 | 0.51 | 0.45 | 0.58 | 0.47 | 1 | - | - | - | - |
| Emotional | 0.20 | 0.22 | 0.09 | 0.28 | 0.17 | 0.28 | 1 | - | - | - |
| Total | 0.67 | 0.48 | 0.40 | 0.63 | 0.72 | 0.72 | 0.67 | 1 | - | - |
| PLIKSi 14 | 0.14 | 0.11 | 0.08 | 0.05 | 0.10 | 0.09 | 0.11 | 0.11 | 1 | - |
| PLIKSi 18 | 0.07 | 0.11 | 0.04 | 0.04 | 0.11 | 0.09 | 0.13 | 0.14 | 0.46 | 1 |

*^AASS: Attention/activity symptoms score; ABS: Awkward behaviours score; TBS: Troublesome behaviours score; SCDC: Social and communication disorder checklist; PLIKSi: Psychosis like symptoms^*

**Table S2. Association of major Y chromosome haplogroups in ALSPAC with the number of** **behavioural traits adjusted for paternal social class and GCSE results**

|  | Effect size of major Y chr haplogroup on behavioural traits:  Beta (95% C.I.)  R haplogroup is the reference | | Adjusted P values |
| --- | --- | --- | --- |
| Behavioural/Psychiatric trait score | **I** | **Other** |  |
| Attention/ Activity symptoms score (DAWBA) | 0.16 (-0.51, 0.83) | 1.36 (0.19, 2.53) | 0.075 |
| Awkward behaviours score (DAWBA) | -0.03 (-0.30, 0.25) | 0.18 (-0.31, 0.66) | 0.81 |
| Troublesome behaviours score (DAWBA) | 0.00 (-0.10, 0.10) | 0.01 (-0.17 , 0.18) | 1.00 |
| SCDC | 0.06 (-0.31, 0.44) | 0.25 (-0.41, 0.90) | 0.74 |
| Hyperactivity traits (SDQ) | 0.06 (-0.17, 0.28) | 0.35 (-0.04, 0.74) | 0.20 |
| Conduct traits (SDQ) | 0.17 (0.03, 0.30) | 0.02 (-0.22, 0.26) | 0.06 |
| Emotional symptoms (SDQ) | 0.04 (-0.11, 0.20) | 0.19 (-0.09, 0.46) | 0.38 |
| Total behavioural traits (SDQ) | 0.12 (-0.33, 0.56) | 0.65 (-0.13, 1.43) | 0.25 |
| PLIKSi age 14 | -0.01 (-0.05, 0.02) | -0.01 (-0.07, 0.06) | 0.76 |
| PLIKSi age 18 | 0.00 (-0.04, 0.03) | 0.01 (-0.05, 0.08) | 0.92 |

*^95% CI: 95% Confidence Interval; DAWBA: Development and Well-Being Assessment; SCDC: Social and communication disorders checklist; SDQ: Strengths and Difficulties Questionnaire; PLIKSi: Psychosis-Like Symptom Interview^*

**Table S3: Association of major mitochondrial DNA haplogroups in ALSPAC with the number of behavioural traits adjusted for maternal and paternal social class and GCSE results**

*^95% CI: 95% Confidence Interval; DAWBA: Development and Well-Being Assessment; SCDC: Social and communication disorders checklist; SDQ: Strengths and Difficulties Questionnaire; PLIKSi: Psychosis-Like Symptom Interview^*

|  | Odds ratios of major mitochondrial DNA haplogroup on behavioural and psychiatric traits: OR (95% C.I.)  HV haplogroup is the reference | | | | | Adjusted P values |
| --- | --- | --- | --- | --- | --- | --- |
| Behavioural/ Psychiatric trait score | **J** | **K** | **TR** | **U** | **Other** |  |
| Attention/ Activity symptoms score (DAWBA) | -0.35  (-1.01, 0.32) | -0.30  (-1.04, 0.44) | 0.03  (-0.62, 0.69) | 0.20  (-0.43, 0.83) | -0.14  (-0.91, 0.64) | 0.78 |
| Awkward behaviours score (DAWBA) | -0.13  (-0.41, 0.16) | -0.06  (-0.38, 0.25) | -0.06  (-0.34, 0.22) | 0.06  (-0.21, 0.32) | -0.08  (-0.41, 0.25) | 0.91 |
| Troublesome behaviours score (DAWBA) | -0.03  (-0.14, 0.07) | -0.02  (-0.13, 0.10) | 0.01  (-0.09, 0.11) | 0.05  (-0.05, 0.15) | 0.06  (-0.06, 0.18) | 0.74 |
| SCDC | -0.26  (-0.63, 0.12) | -0.25  (-0.66, 0.17) | -0.20  (-0.57, 0.17) | 0.05  (-0.30, 0.40) | -0.22  (-0.66, 0.22) | 0.50 |
| Hyperactivity traits (SDQ) | -0.03  (-0.26, 0.20) | 0.05  (-0.20, 0.31) | 0.21  (-0.02, 0.45) | 0.05  (-0.17, 0.26) | 0.06  (-0.21, 0.33) | 0.59 |
| Conduct traits (SDQ) | 0.08  (-0.06, 0.23) | -0.07  (-0.23, 0.09) | 0.09  (-0.06, 0.23) | 0.03  (-0.11, 0.16) | -0.02  (-0.20, 0.15) | 0.58 |
| Emotional symptoms (SDQ) | 0.08  (-0.09, 0.26) | -0.10  (-0.29, 0.09) | -0.07  (-0.24, 0.11) | 0.03  (-0.13, 0.20) | -0.16  (-0.36, 0.05) | 0.35 |
| Total behavioural traits (SDQ) | 0.18  (-0.30, 0.65) | -0.03  (-0.55, 0.48) | 0.23  (-0.24, 0.70) | 0.14  (-0.30, 0.58) | -0.26  (-0.81, 0.29) | 0.70 |
| PLIKSi age 14 | 0.03  (-0.08, 0.14) | -0.07  (-0.19, 0.04 | 0.04  (-0.06, 0.15) | -0.07  (-0.17, 0.03) | -0.02  (-0.15, 0.10) | 0.37 |
| PLIKSi age 18 | -0.02  (-0.15, 0.11) | 0.02  (-0.11, 0.16) | 0.07  (-0.05, 0.20) | -0.02  (-0.13, 0.10) | -0.14  (-0.29, 0.01) | 0.29 |

**Table S4. Odds ratios of Y chromosome haplogroups subgroups on binary psychiatric trait measures from logistic regression adjusted for paternal social class and GCSE results**

| Behavioural/ Psychiatric trait score | P Values | | | |
| --- | --- | --- | --- | --- |
|  | **N** | **Unadjusted** | **N** | **Adjusted** |
| Attention/ Activity symptoms score (DAWBA) | 3265 | 0.26 | 2400 | 0.25 |
| Awkward behaviours score (DAWBA) | 3251 | 0.67 | 2390 | 0.48 |
| Troublesome behaviours score (DAWBA) | 3262 | 0.29 | 2396 | 0.58 |
| SCDC | 3253 | 0.39 | 2392 | 0.16 |
| Hyperactivity traits (SDQ) | 3300 | 0.64 | 2461 | 0.63 |
| Conduct traits (SDQ) | 3304 | 0.76 | 2460 | 0.43 |
| Emotional symptoms (SDQ) | 3301 | 0.06 | 2460 | 0.09> 0.006^1^ |
| Total behavioural traits (SDQ) | 3298 | 0.74 | 2457 | 0.78 |
| PLIKSi age 14 | 2859 | 0.95 | 2059 | 0.98 |
| PLIKSi age 18 | 1749 | 0.74 | 1228 | 0.56 |

*^95% CI: 95% Confidence Interval; DAWBA: Development and Well-Being Assessment; SCDC: Social and communication disorders checklist; SDQ: Strengths and Difficulties Questionnaire; PLIKSi: Psychosis-Like Symptom Interview^*

*^1 Smallest adjusted p value compared to multiple testing threshold^*

**Table S5. Association of Y chromosome haplogroup subgroups in ALSPAC with the number of behavioural traits adjusted for paternal social class and GCSE results**

| Behavioural/ Psychiatric trait score | P Values | | | |
| --- | --- | --- | --- | --- |
|  | **N** | **Unadjusted** | **N** | **Adjusted** |
| Attention/ Activity symptoms score (DAWBA) | 3265 | 0.40 | 2400 | 0.07 |
| Awkward behaviours score (DAWBA) | 3251 | 0.71 | 2390 | 0.22 |
| Troublesome behaviours score (DAWBA) | 3262 | 0.78 | 2396 | 0.59 |
| SCDC | 3253 | 0.39 | 2397 | 0.05>0.006^2^ |
| Hyperactivity traits (SDQ) | 3300 | 0.64 | 2461 | 0.63 |
| Conduct traits (SDQ) | 3304 | 0.61 | 2462 | 0.26 |
| Emotional symptoms (SDQ) | 3301 | 0.14 | 2462 | 0.20 |
| Total behavioural traits (SDQ) | 3298 | 0.76 | 2459 | 0.29 |
| PLIKSi age 14 | 2859 | 0.68 | 2062 | 0.53 |
| PLIKSi age 18 | 1749 | 0.11 | 1270 | 0.13 |

*^95% CI: 95% Confidence Interval; DAWBA: Development and Well-Being Assessment; SCDC: Social and communication disorders checklist; SDQ: Strengths and Difficulties Questionnaire; PLIKSi: Psychosis-Like Symptom Interview^*

*^1 Smallest adjusted p value compared to multiple testing threshold^*

**Table S6. Odds ratios of mitochondrial DNA chromosome haplogroup subgroups on binary behavioural trait measures from logistic regression adjusted for paternal social class and GCSE results**

| Behavioural/ Psychiatric trait score | P Values | | | |
| --- | --- | --- | --- | --- |
|  | **N** | **Unadjusted** | **N** | **Adjusted** |
| Attention/ Activity symptoms score (DAWBA) | 5674 | 0.95 | 3857 | 0.88 |
| Awkward behaviours score (DAWBA) | 5646 | 0.20 | 3841 | 0.66 |
| Troublesome behaviours score (DAWBA) | 5649 | 0.90 | 3843 | 0.83 |
| SCDC | 5651 | 0.19 | 3850 | 0.21 |
| Hyperactivity traits (SDQ) | 5731 | 0.71 | 3926 | 0.90 |
| Conduct traits (SDQ) | 5738 | 0.90 | 3925 | 0.58 |
| Emotional symptoms (SDQ) | 5731 | 0.23 | 3927 | 0.12> 0.006^1^ |
| Total behavioural traits (SDQ) | 5723 | 0.77 | 3922 | 0.35 |
| PLIKSi age 14 | 5154 | 0.44 | 3426 | 0.42 |
| PLIKSi age 18 | 3516 | 0.18 | 2369 | 0.63 |

*^95% CI: 95% Confidence Interval; DAWBA: Development and Well-Being Assessment; SCDC: Social and communication disorders checklist; SDQ: Strengths and Difficulties Questionnaire; PLIKSi: Psychosis-Like Symptom Interview^*

*^1 Smallest adjusted p value compared to multiple testing threshold^*

**Table S7. Association of mitochondrial DNA haplogroup subgroups in ALSPAC with the number of behavioural traits adjusted for maternal and paternal social classes and GCSE results**

| Behavioural/ Psychiatric trait score | P Values | | | |
| --- | --- | --- | --- | --- |
|  | **N** | **Unadjusted** | **N** | **Adjusted** |
| Attention/ Activity symptoms score (DAWBA) | 5674 | 0.65 | 3847 | 0.99 |
| Awkward behaviours score (DAWBA) | 5646 | 0.28 | 3841 | 0.53 |
| Troublesome behaviours score (DAWBA) | 5649 | 0.85 | 3843 | 0.98 |
| SCDC | 5651 | 0.11 | 3850 | 0.16 |
| Hyperactivity traits (SDQ) | 5731 | 0.71 | 3926 | 0.90 |
| Conduct traits (SDQ) | 5738 | 1.00 | 3929 | 0.74 |
| Emotional symptoms (SDQ) | 5731 | 0.18 | 3927 | 0.25 |
| Total behavioural traits (SDQ) | 5723 | 0.95 | 3922 | 0.90 |
| PLIKSi age 14 | 5154 | 0.32 | 3429 | 0.23 |
| PLIKSi age 18 | 3516 | 0.03 | 2370 | 0.01> 0.006^1^ |

*^95% CI: 95% Confidence Interval; DAWBA: Development and Well-Being Assessment; SCDC: Social and communication disorders checklist; SDQ: Strengths and Difficulties Questionnaire; PLIKSi: Psychosis-Like Symptom Interview^*

*^1 Smallest adjusted p value compared to multiple testing threshold^*
